# Augment Single-cell RNA-seq data with Surface Protein Levels using Gene set-based Deep Learning and Transfer Learning Methods

**DOI:** 10.1101/2024.04.29.591655

**Authors:** Md Musaddaqul Hasib, Tinghe Zhang, Jianqiu Zhang, Shou-jiang Gao, Yufei Huang

## Abstract

As scRNA-seq becomes increasingly accessible, providing a cost-efficient method to augment surface protein levels from gene expression measurements are desirable. We proposed a machine learning approach that includes a novel geneset neural network (GS-NN) that aims to learn robust and biologically meaningful features and a highly efficient transfer learning strategy to address cross-dataset differences. We conducted comprehensive experiments to show the improvements of the proposed methods. Specifically, we demonstrate that GS-NN learns more robust features to achieve better cross-subject performance than other machine learning approaches. Transfer learning further improves that of GS-NN by reducing dataset differences through highly efficient fine-tuning. The unique genesets design of GS-NN also allows identification of functions contributing to the prediction and improvement of the proposed strategy. Overall, this study reports a novel approach to robustly augment.

**Key Points:**

- The article presents a machine learning approach, Geneset Neural Network(GS-NN) to augment surface protein levels from single-cell RNA sequencing(scRNA-seq) gene expression data.
- The GS-NN aims to learn robust and biologically meaningful features, and the approach includes a highly efficient transfer learning strategy to address cross-dataset differences in scRNA-seq data.
- Comprehensive experiments demonstrate that GS-NN learns more robust features using trasfer learning techniques achieving better cross-subject performance compared to other machine learning approaches.
- The unique geneset-based architecture of GS-NN allows the identification and interpretion of biological functions contributing to the prediction of cell surface protein level.
- GS-NN’s architecture is conveniently transferrable across datasets, making it valuable tool for researchers working with diverse scRNA-seq datasets.

## 1. Introduction

Gene expressions within a cell are tightly linked to the level of surface proteins. The modulation of gene expression can occur due to the impact of proteins on RNA. It is recognized that proteins are highly abundant compared to RNA and functionally directly involved in cell-cell interactions and signaling[1]. Despite having immense importance of quantified proteins level information, most projects including Human Cell Atlas project[2] only quantify transcriptome considering technological barriers. Recently developed, CITE-seq[3]multi-omics technology measures RNA and protein expression simultaneously in single cell resolution. This combination of transcriptomics and proteomics data in CITE-seq hold potential to provide better functional insights into cellular responses, signaling pathways, and interactions within complex biological system that might have been missed by scRNA-seq. However, this new technology is not as cost effective as scRNA-seq yet. Utilizing the existing CITE-seq data to augment surface protein levels with machine learning in the vast amount existing scRNA-seq data would significantly expand the understanding of cell states and functions and therefore highly valuable. This paper considers the problem of predicting surface protein levels using expression profiles from scRNA-seq. Particularly, we focus on the more realistic setting of cross subjects/datasets prediction, i.e., making predictions in new datasets different from the training datasets. Biological data inherit cross dataset variations known as batch effects. Batch effects are the systematic non-biological differences between batches and inevitable because data are often generated at different times and the batches are confounding biological variations. Due to this effect, a well-trained model on a dataset performs better within that dataset but could perform poorly when tested for cross datasets. cTP-net[4], a deep learning-based approach, demonstrates this phenomenon while predicting surface protein levels from gene expressions using CITE-seq PBMC and CBMC data. Their model is built using only the fully connected neural network that is prone to overfit the data and thus did not work well for cross dataset predictions. An ensemble machine learning approach was proposed to improve the accuracy of predicting single cell protein abundance without using any deep learning[5]. They selected only the top 20 highly correlated genes to train the ensemble model and predict surface protein within the dataset. They did not evaluate the cross-data performance as cPT-net did. This approach ignores a lot of expressed genes in a cell that may have regulatory effects on surface protein expression. Therefore, it would not be able to reveal the contributions of those regulatory genes.

In this paper, we aim to develop a deep transfer learning model that is more robust and biologically inspired. In scRNA-seq analysis, applying a batch correction method such as [6], BBKNN[7], etc to mitigate batch effects is a popular strategy. However, besides added computations, special care needs to be taken when using this strategy to a specific dataset to balance between overly correcting real biological differences and properly removing batch effects. Instead, we investigate transfer learning, a widely adopted strategy to address domain and even task differences in natural language processing and computer vision. In addition, we aim to improve the design of deep learning models such that while predicting surface protein levels from gene expression also facilitate meaningful, easier functional interpretation. Direct adaptation of popular DL models such as convolution neural networks (CNNs)[8] in this task may not be suitable because CNNs exploit pixel/text correlations within a local region, they cannot fully capture functional gene interactions in expression data. The gene expression data is unordered (Unlike pixels in images) where interacting genes might be separated in the unordered expression vector. We propose a gene-set-based deep learning ensemble approach (GS-NN) that takes gene expressions of biological pathways (functions) as input and optimizes for surface protein level prediction task and then integrates/ensembles those gene-set embedding to do final prediction. The process of selecting gene-sets provides an important avenue for investigators to directly inject their state of understanding and hypotheses about the pathways into GS-NN so that GS-NN’s prediction and functional interpretation are attuned to the study’s goal. Selecting gene-sets has some advantages over selecting genes in the DL models. Gene-sets usually have a small number of genes compared to whole transcriptomics (>20K genes) and learning gene-gene correlation within a gene-set is less prone to overfitting. Instead of removing standalone genes we chose to remove poorly performed gene-sets from GS-NN since genes within gene-sets act together and are biologically more meaningful. We also argue that GS-NN learns more robust features of gene sets and thus transferring knowledge reduces the batch effect.

The scripts and data used to perform the main analyses are available in zenodo https://doi.org/10.5281/zenodo.10447725

## 2. Materials and Methods

### 2.1.1. Datasets

We downloaded the human non-small cell lung cancer (NSCLC) [9]CITE-seq datasets. For this study, we used 10 datasets including 5 from normal lung tissues and 5 from lung tumor tissues mixed with multiple subjects (Table 1). Each dataset contains a total of 21 surface protein level abundance. The expression count data of filtered cells are public in a GitHub repository. All human sequencing data is available on NCBI with ‘BioProject’ ID PRJNA609924 and GEO accession GSE154826.

**Table 1.**
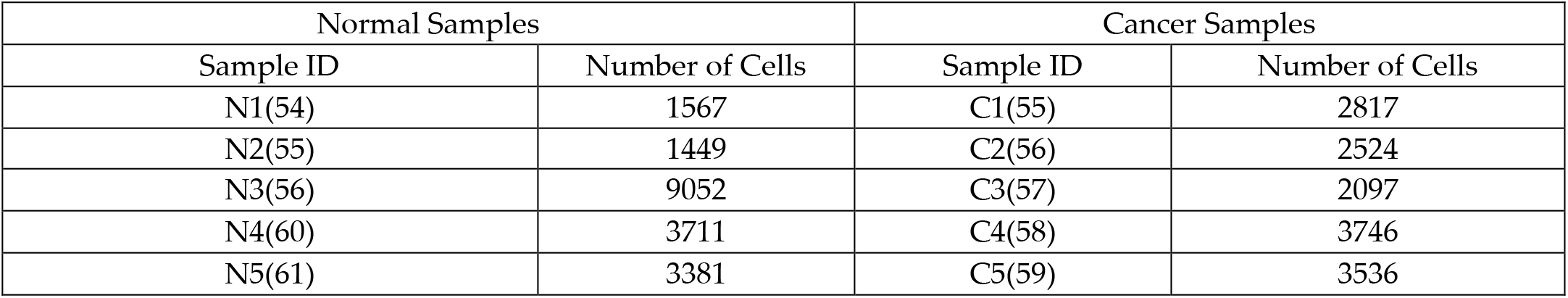
Human non-small cell lung cancer CITE-seq datasets summary.

**Surface proteins:** ‘CD11b’, ‘CD11c’, ‘CD123’, ‘CD14’, ‘CD141’, ‘CD16’, ‘CD19’, ‘CD1c’,’CD206’, ‘CD24’, ‘CD27’, ‘CD3’, ‘CD33’, ‘CD38’, ‘CD4’, ‘CD56’, ‘CD62L’, ‘CD64’, ‘CD66b’, ‘CD8’, ‘HLA-DR’

### 2.1.2. Imputation of gene expression data and normalization of surface protein levels

We first denoised the gene expression and normalized surface protein abundance as a preprocessing step. The scRNA-seq count matrix was denoised using the affinity matrix-based MAGIC [10]algorithm, which produces more accurate estimates of each cell’s RNA transcript relative abundance. Compared to the raw counts, the denoised relative expression values have significantly improved correlations with their corresponding protein measurements. Surface protein levels were also normalized using the scanpy package [11]. We normalized the protein abundance of each cell by the total abundance of all proteins, so every cell has the same total protein abundance after normalization.

#### 3. Geneset selection

Total 186 canonical pathways genesets derived from the KEGG pathway database were obtained from the MSigDB database[12] publicly available for download. On top of that, we also added 7 custom cell type gene-sets obtained from the Cell Match database[13].

### 2.2. Methods

Here, we propose a novel gene-set-based ensemble neural network (GS-NN) for gene expression-based prediction of surface protein levels. A transfer learning approach based on GS-NN is further implemented for a cross-dataset prediction that can reduce the inherited batch effects.

#### 2.2.1. Proposed geneset neural network (GS-NN)

Given a CITE-seq dataset that contains imputed and normalized gene expressions *X* ∈ *R*^*C*×*G*^ and normalized surface protein levels *Y* ∈ *R*^*C*×*P*^, where *C* is the number of cells, *G* is the total number of genes, and *P* is the number of surface proteins. The goal is to train a GS-NN model to predict *Y* from *X* (Fig. 1). To this end, a set of gene-sets is first obtained. Then, for geneset *j*, an NN *f*_*j*_ (θ_*j*_, *X*_*j*,_ *Y*), is trained to predict *Y* using only the expression data from the genes of that geneset or *X*_*j*_, where θ_*j*_ are weights learned during training. Afterward, we can select *M* highest performing geneset models to construct GS-NN. To this end, a linear ensemble layer is applied to integrate the output features of each geneset NN’s pre-activations (Fig. 1). We can add additional layers to further reduce the dimension of features before feeding them to the regression layer to predict *Y*. While training GS-NN, weights of geneset NNs can be fixed or fine-tuned. In this study, we choose to freeze the weights of individual geneset NNs to keep features that already learnt and train only additional layers after ensemble. The final GS-NN can be expressed as *Y* = *F* (θ, *X*), where θ represents all weights including pre-trained weights from geneset NN’s.

**Figure 1:**
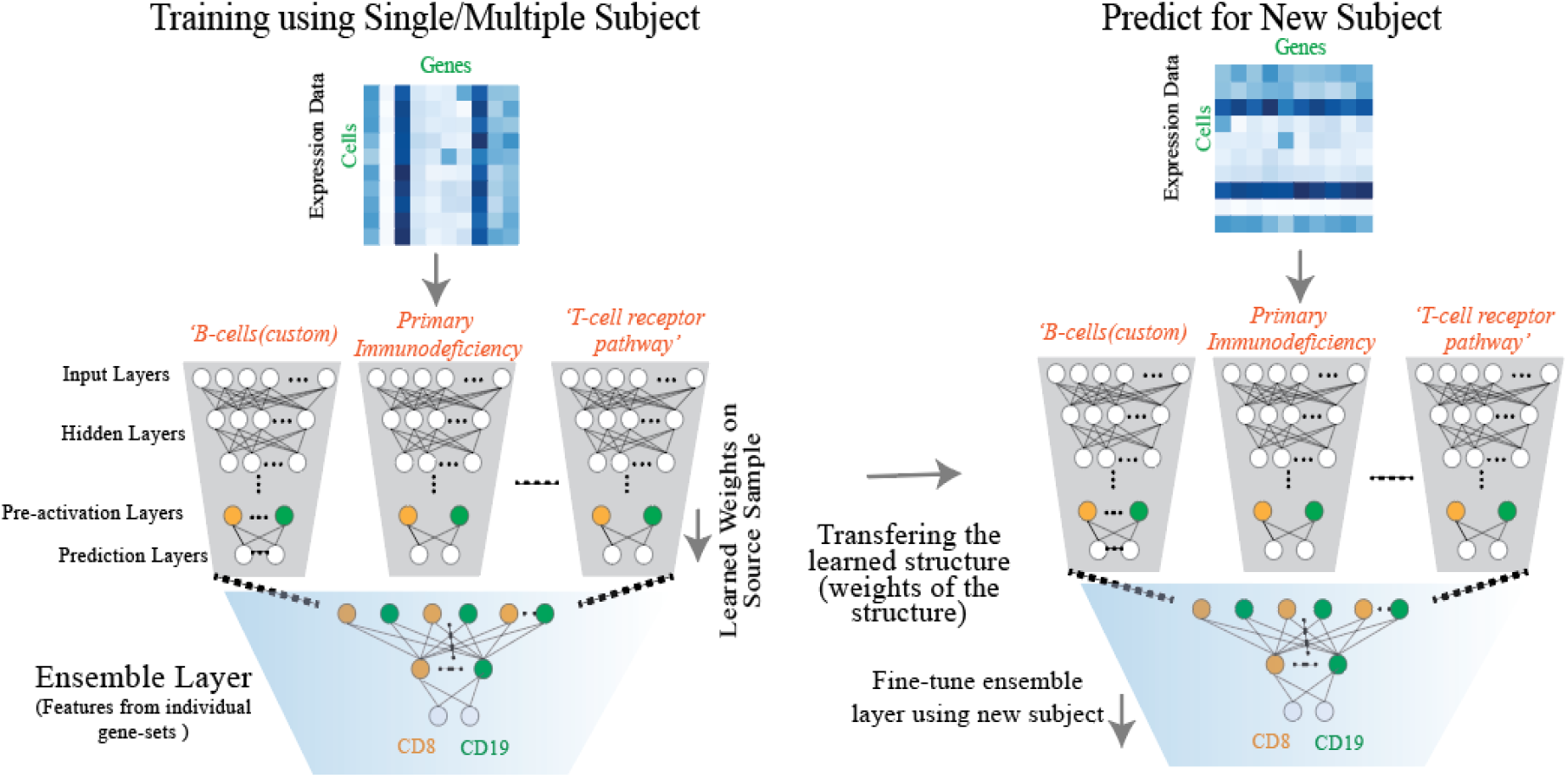
Predicting cross-subject cell surface protein level prediction using proposed Geneset Neural Network (GS-NN) and Transfer Learning

#### 2.2.2. Transfer learning with GS-NN for cross-dataset prediction

For cross-dataset prediction using GS-NN, we propose a transfer learning strategy. Suppose that we have a source dataset, from which we train a GS-NN model. The goal is to adapt the trained GS-NN to a new target dataset, where there are limited training samples. With transfer learning, we fine-tune the GS-NN using the target training data. Applying fine-tuning help source GS-NN to adapt to features of the target dataset. GS-NN has an inherent advantage of architecture design because GS-NN are pre-trained independently and can be easily transferred to a new dataset and even for a new task. In this study, we fine-tune only the ensemble layers much faster than training from scratch using a new dataset.

#### 2.2.3. Model training

All ten samples (Normal and Cancer) were first divided into 3-fold and 1-fold was separated for testing. This training and testing data were consistent throughout all experiments and for all algorithms. For the performance evaluation, we used the Pearson correlation coefficient (PCC) between predicted surface protein level and ground truth surface protein level and then averaged them across all proteins.

Building GS-NN requires integrating geneset NNs models optimized for protein level prediction tasks. Thus, as the first step, we trained total of 194 geneset models, one for a geneset, using Auto-Keras[14]. Auto-keras automatically finds the best-optimized architecture for the prediction. There are a few hyperparameters that we selected. We set ‘block_type= vanilla’, which allowed the search within standard convolutional neural networks, ‘normalze= True’, which normalized each training batch. The ‘batch_size=32’, ‘max_trials = 5’ which tried 5 models, each optimized for different architectures, ‘epoch = 50’ is the number passes of the training dataset. Auto-Keras returned the best performed model out of these 5 models it tried. Then, we select some gene-sets using the PCC performance threshold to build GS-NN. To integrate gene-set NNs we use a linear stacking ensemble layer followed by a dense layer using 50 neurons and a regression layer to do the final prediction of surface protein levels.

We compared the performance of GS-NN with other machine learning models as baseline including support vector regression (SVR)[15], decision tree regression (DT)[16], gradient boost regression (GB)[17] and vanilla CNN[8]. Because these baseline models are not gene-set-based, we selected 500 highly variable genes as input. We used different kernels including “linear” and “radial bias function (rbf)” SVR, different depth for decision DT regressor, number of estimators and maximum depth parameters to maximize prediction performance. Vanilla-CNN is optimized using Auto-Keras as explained in the previous paragraph.

### 2.3. Interpretation of GS-NN

The purpose of the interpretation of GS-NN is to investigate how GS-NN makes predictions of surface protein levels. By interpretation, we can reveal gene-sets important for surface protein level predictions.

#### 2.3.1. Identifying marker genes for surface protein prediction using integrated gradients (IG)

IG[18] is a feature attribution-based technique that explains the relationship between a model’s predictions and features and assigns an attribution score to each feature for specific input. It has become a popular interpretability technique due to its broad applicability to any differentiable model and efficient implementation. IG computes attribution scores by accumulating gradients samples interpolated between the baseline value and the current input. Given an input gene expression of a cell *x*_*k*_ to GS-NN function *F* where, *k* is the cell index, IG computes an attribution score for each input gene *ϕ*_*kjl*_ (*F, x*_*k*_), where *l* is a gene ID. A significant positive or negative attribution score of a gene indicates that the gene significantly impacts the GS-NN output *F* (*x*_*k*_). An attribution scores close to zero indicates that the gene does not influence *F* (*x*_*i*_).

#### 2.3.2. Functional enrichment analysis using Fisher exact test

Fisher’s exact test[19] is a classic statistical test to perform the functional enrichment analysis that identifies gene-sets that share an unusually large number of genes with the rank list of genes using attributions scores derived from IG. Our two-sided Fisher’stest outputs two values, the odd ratio that gives us a quantitative measurement of genes from a geneset that appears on the top of the ranked list and the p-value that determines how significant the test is.

## 3. Results

Here in the result, we discuss the performance of GS-NN for the prediction of cell surface protein level using gene expression data. We first consider the within-dataset prediction in normal and cancer samples separately and compare the proposed GS-NN model with multiple machine learning models. Then, we investigate the direct cross-dataset prediction performance of GS-NN, where we explore the performance of GS-NN that is trained with a single dataset and multiple datasets. Next, we investigate the transfer learning performance of GS-NN for two cross-dataset prediction scenarios and strategies including transferring from one dataset to another holdout dataset and transferring from multiple datasets to one holdout dataset.

### 3.1.1. Investigating the surface protein levels prediction performance of GS-NN within dataset using scRNA-seq gene expression

This section discusses the within dataset performance, i.e., training and test samples are from the same dataset; thus no cross-dataset differences. We selected one dataset from normal tissue (N5) and one from tumor dataset (C5) to build GS-NN and test its performance. They are also the test dataset for transfer learning discussed in later sections. After training geneset NNs, we sorted the geneset models based on their average PCCs.

For normal sample N5, we found that the custom-built ‘B-cells’ gene-set achieved the highest average PCC(0.73) and ‘*CYTOKINE_CYTOKINE_RECEPTOR_INTERACTION’* ranks the second (PCC:0.72) (Table 2). They are the top two gene-sets for cancer sample C5 as well. These top gene-sets are closely connected to extracellular proteins and immune responses. For example, the ‘B-cells’ geneset includes B cell marker genes. ‘Cytokine’ are crucial intercellular regulators related to innate and adaptive immune responses. Nevertheless, we observe that the top genesets for C5 have 0.04 points lower performance than those in N5. This result suggests that tumor sample C5’s gene expressions to their protein levels have more considerable variations than N5.

**Table 2.** Top-ranked. genesets based on performance of predicting cell surface protein level using CNN.

| Top Genesets for Normal Sample (N5) | Average Correlation | Top Gene-sets for Cancer Sample(C5) | Average Correlation |
| --- | --- | --- | --- |
| B_cells (Custom) | 0.73 | CYTOKINE_CYTOKINE_RECEPTOR_INTERACTION | 0.691 |
| CYTOKINE_CYTOKINE_RECEPTOR_INTERACTION | 0.72 | B_cells (Custom) | 0.691 |
| PRIMARY_IMMUNODEFICIENCY | 0.72 | TYPE_I_DIABETES_MELLITUS | 0.689 |
| TOLL_LIKE_RECEPTOR_SIGNALING_PATHWAY | 0.71 | GRAFT_VERSUS_HOST_DISEASE | 0.678 |
| Monocytes (Custom) | 0.71 | ALLOGRAFT_REJECTION | 0.676 |

If we sort the gene-sets based on their PCC performance for predicting CD19, which is a B-cell marker, we see that ‘PRIMARY_IMMUNODEFICIENCY’ and ‘B_cells’ are the top two genesets with a correlation of 0.83 and 0.82, respectively for normal sample (N5) (Table 3). ‘B_CELL_RECEPTOR_SIGNALING_PATHWAY’ ranked fifth with 0.79 PCC for the same sample. Higher correlations are observed for the cancer sample than for the normal sample. This might be because there is a higher immune response in tumor. ‘CYTOKINE_CYTOKINE_RECEPTOR_INTERACTION’, ‘Protein genes’ and ‘B-cells’ with 0.89. We observe that for the geneset model ‘B-cells’, CD19 PCC performance is significantly higher than the average PCC performance in N5 and C5. Because CD19 is a B cells marker, this result suggests that individual gene-set models can indeed capture surface protein-related features.

**Table 3.** Top-ranked. genesets based on performance of predicting B-cell surface protein marker CD19 using CNN.

| Normal sample N5(CD19) |  | Cancer Sample C5 (CD19) |  |
| --- | --- | --- | --- |
| Gene-set | Correlation | Gene-set | Correlation |
| PRIMARY_IMMUNODEFICIENCY | 0.837 | CYTOKINE_CYTOKINE_RECEPTOR_INTERACTION | 0.89 |
| B_cells (Custom) | 0.821 | Protein genes (Custom) | 0.89 |
| TYPE_I_DIABETES_MELLITUS | 0.812 | B_cells(Custom) | 0.85 |
| LEISHMANIA_INFECTION | 0.804 | Monocytes (Custom) | 0.85 |
| PROSTATE_CANCER | 0.803 | PRIMARY_IMMUNODEFICIENCY | 0.83 |
| B_CELL_RECEPTOR_SIGNALING_PATHWAY | 0.799 | LEISHMANIA_INFECTION | 0.83 |

To build GS-NN, for normal sample N5, we selected 17 genesets using PCC threshold 0.70 for cancer dataset C5, we selected 9 genesets using PCC threshold 0.67. Then, we trained the ensemble layer to combine these geneset models for each dataset. We found that these number of gene-sets are sufficient to maximixe surface protein prediction performance. When we compare the average PCC performance of GS-NN with other predictors, GS-NN becomes one of the best predictors for both N5 and C5 (Figure 2). Besides the average PCC performance, we also examine individual protein prediction performances. We see that PCC performance for predicting Natural killer (NK) cell marker CD16 is 0.02 points better for N5 (Figure 2-A) and 0.04 points better for C5 than GB(Figure 2-B). Predicting B-cell marker CD19 shows 0.05 points better than GB for N5(Figure 2-A). HLA-DR is a marker for monocytes, B-cells and one of the top predicted proteins using GS-NN, vanilla CNN and GB. These results show that GS-NN is one of the best predictors for within dataset prediction.

**Figure 2.**
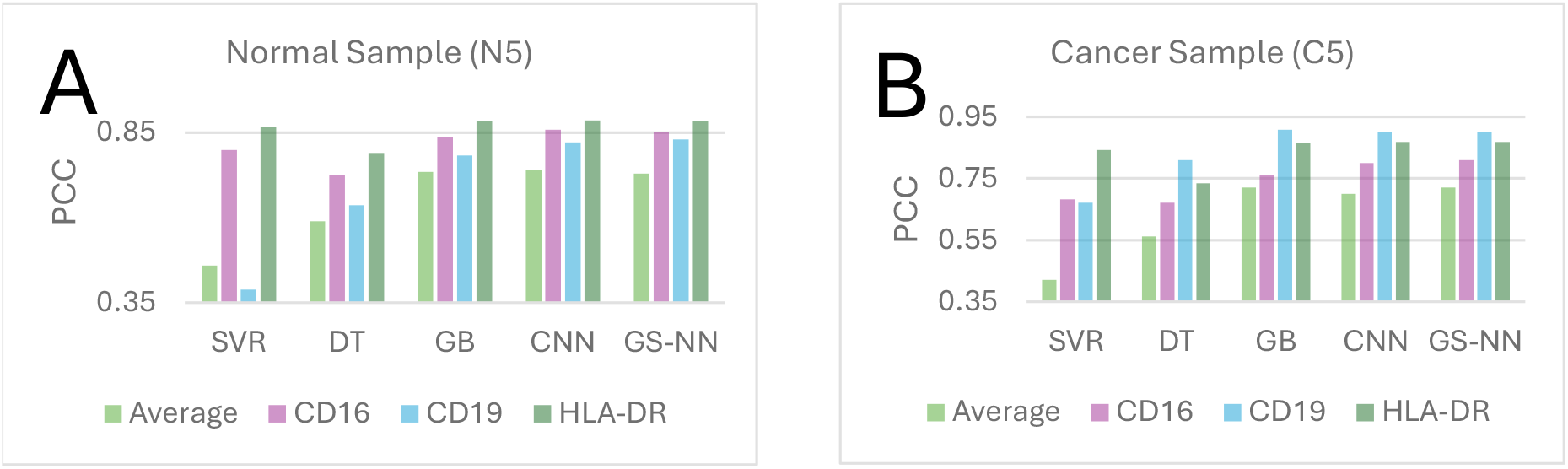
Within-subject prediction performance of cell surface protein level using various traditional machine learning and Deep learning methods.

### 3.1.2. Investigating direct cross-subject prediction performance of cell surface protein level

In the previous section, we summarized the results of GS-NN performance by comparing other predictors using N5 and C5 datasets separately. This section shows the performance of applying GS-NN trained in N5 or C5 to make predictions in the remaining 4 normal or cancer datasets. First, among normal datasets, we trained GS-NN with 2 folds of N5 samples and tested with 1-fold from each of N1, N2, N3 and N4. Similarly, for cancer datasets, we trained GS-NN with 2 folds of the C5 dataset and tested with 1-fold from each of C1, C2, C3 and C4, separately. We keep it consistent with the testing setting of N5 and C5 because we are going to test their transferability to other datasets in a later section. We see comparable performance between GS-NN and CNN for normal samples, which are better than other models (Fig. 3). Among the 4 samples, there is considerable variation in performance with N4 having the highest (0.71) and N3 the lowest (0.54) for GS-NN (Fig. 3-A). This also suggests that N5 has fewer differences with N4 compared to other samples. In contrast to normal samples, GS-NN significantly outperforms other models for cancer samples, improving average of 0.30 and 0.15 points in PCC over CNN and GB, respectively (Fig.3B) We see fewer variations and higher PCCs in performance among individual samples compared to normal samples, suggesting there are less differences between cancer samples than between normal samples. Fig.3C shows GS-NN has significantly higher cross-subject average PCC for both normal and cancer subjects. In conclusion, the improved direct cross-dataset performances by GS-NN especially for cancer samples show that GS-NN learned more robust features than CNN and other models.

**Figure 3.**
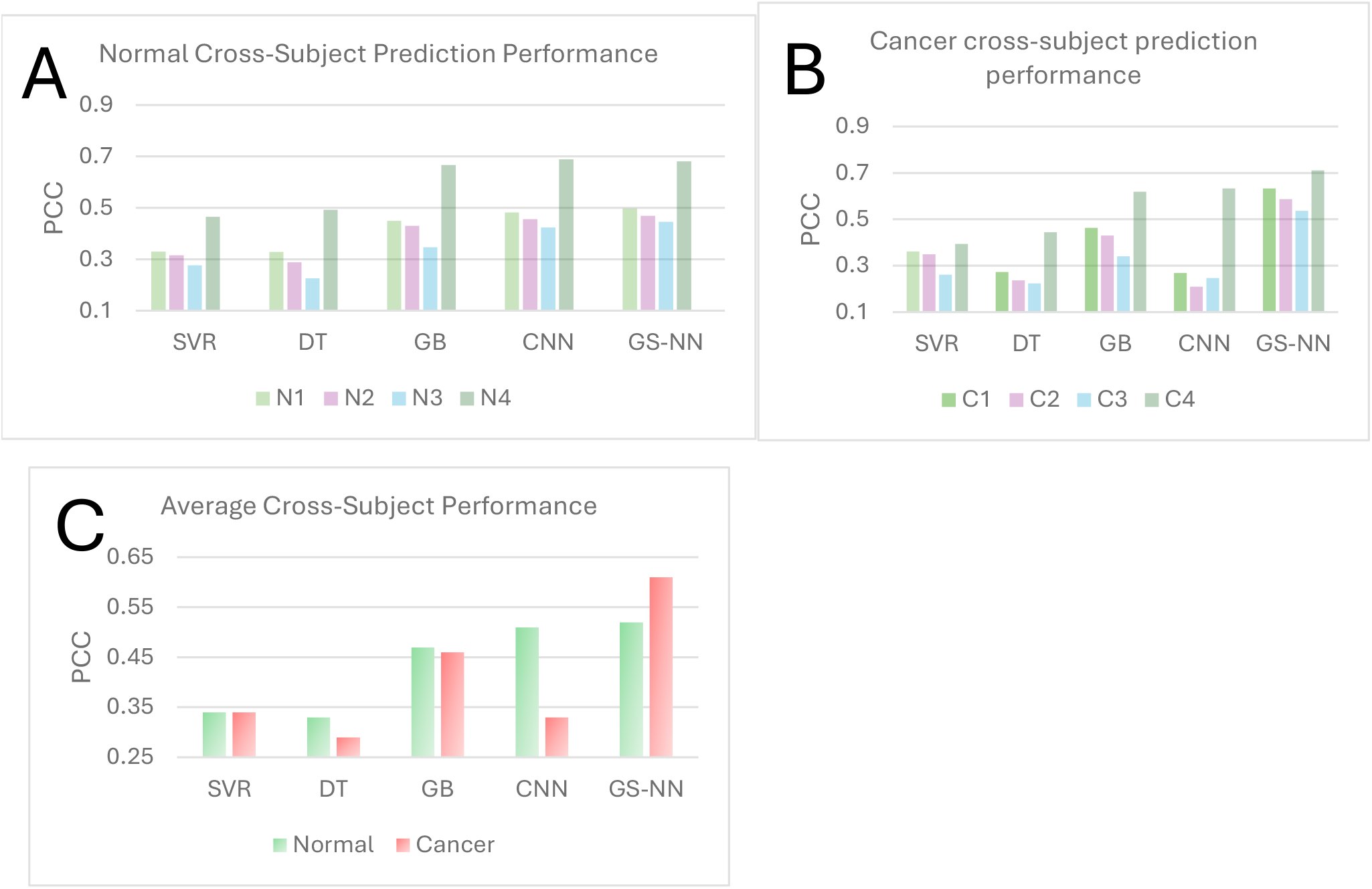
Cross-subject prediction performance of cell surface protein level using various traditional machine learning and Deep learning methods.

Next, we examine if increasing the training sample size allows GS-NN to learn more robust features and thus improves the performance of direct cross-dataset prediction. To do that, we combined four datasets (N1, N2, N4, N5) for normal samples and took 2-folds from each dataset to train the models and then we tested them with 1-fold samples from N3. We do the same for cancer sample C3. We choose N3 and C3 because they are associated with the lowest performance (Fig. 3). We observe that GS-NN still outperforms other machine learning models for both N3 and C3 (Fig. 4). Compared to the performances using only N5 as the training dataset in (Fig. 3), we see an improvement from 0.446 to 0.46 for N3 (Fig. 3A) but decreased performance for C3 (Fig. 3B). This may suggest that there is significantly more variation across cancer samples and combining them weakens instead of strengthening the robust features.

**Figure 4.**
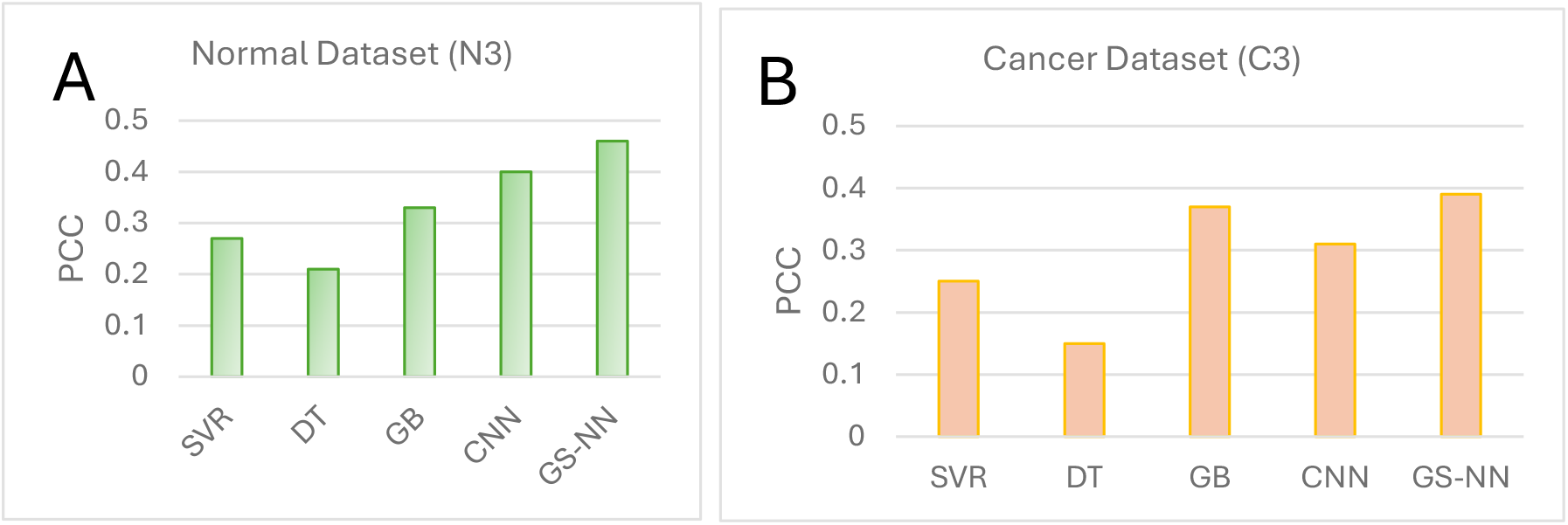
Cross-Subject cell surface prediction performance training after combining multiple subjects.

### 3.1.3. Transferring learning using GS-NN can improve surface protein level prediction performance

Next, we examine the performance of transfer learning with GS-NN. In the previous section, we found that normal sample N3 cancer sample C3 has the lowest cross-sample performance. Now we examine whether it is possible to improve the performance using transfer learning. Similarly, we use 1-fold of N3 or C3 as the test data but different from direct cross-dataset prediction, we fine-tune only the ensemble layer of GS-NN on the remaining 2 folds before making predictions. We consider transferring from N5 or C5, to be comparable to the results in Section 3.1.2. As a baseline, we also train GS-NNs using only 2-fold N3 or C3, i.e., the training has no access to other samples except N3 or C3. We see that fine-tuning GS-NN from N5 or C5 improves about 0.03 points over the baseline GS-NN (Fig. 5). This result demonstrates the effectiveness of transfer learning. We further investigate whether transferring a model trained on more datasets can improve performance. We take the models trained in Section 3.1.2 using 4 normal or cancer samples and perform the fine-tuning on N3. Now, consistent with the results in Fig. 4-B, we see an improvement in transferring from a large dataset only for normal but not cancer samples (Fig.5-B), which implies again more sample variation in cancer samples.

**Figure5.**
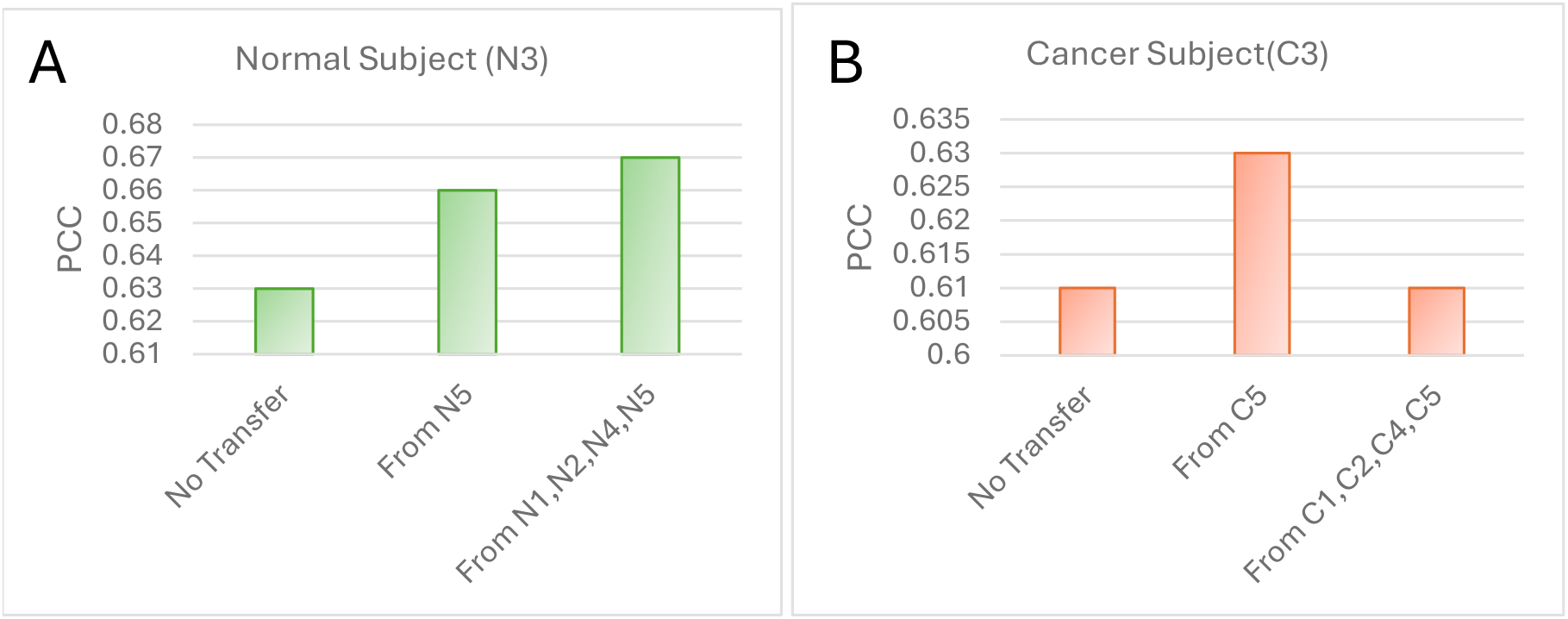
Transfer learning performance of cross dataset using GS-NN.

### 3.2. Functional interpretation of cell surface proteins using GS-NN using transfer learning

In this section, we investigate the functional relevance for predicting cell surface proteins using gene expression data revealed by GS-NN and examine the changes at the functional level that are accompanied with the improved performance by transfer learning. First, we compute the average attribution scores across test samples for each input gene to obtain their global contribution to the prediction. Then, we rank them based on global attribution scores and performed enriched of the top 20% genes in the genesets that GS-NN is built on using the Fisher’s exact test. Here, we focus on significantly enriched genesets (p-adjusted <=0.05) for both normal and cancer subject, predicting B-cell marker CD19 under different transfer learning setups.

First, we investigate functional difference with and without transfer learning using one dataset for normal and cancer sample. For without transfer, we directly predict 1-fold data from N3 using 2 fold N5 GS-NN for normal, for cancer we directly predict 1-fold data from C3 using 2 fold C5 GS-NN. In contrast, for with transfer, we fine-tune GS-NN using N5 with 2 fold data from N3 for normal and cancer similarly. The CD19 PCC perforamnce for N3 improved from 0.81 without transfer to 0.83 after transfer from N5. We find 10 enriched genesets with tranfering and 9 without transfering (Fig. 6A). Immune related geneset ‘CYTOKINE_CYTOKINE_RECEPTOR_INTERACTION’ become the top enriched pathway. Another immune related geneset ‘WNT_SIGNALING_PATHWAY’ only enriched after fine tuning. This suggests, transfer from N5 GS-NN focus on immune-related genesets.

**Figure 6.**
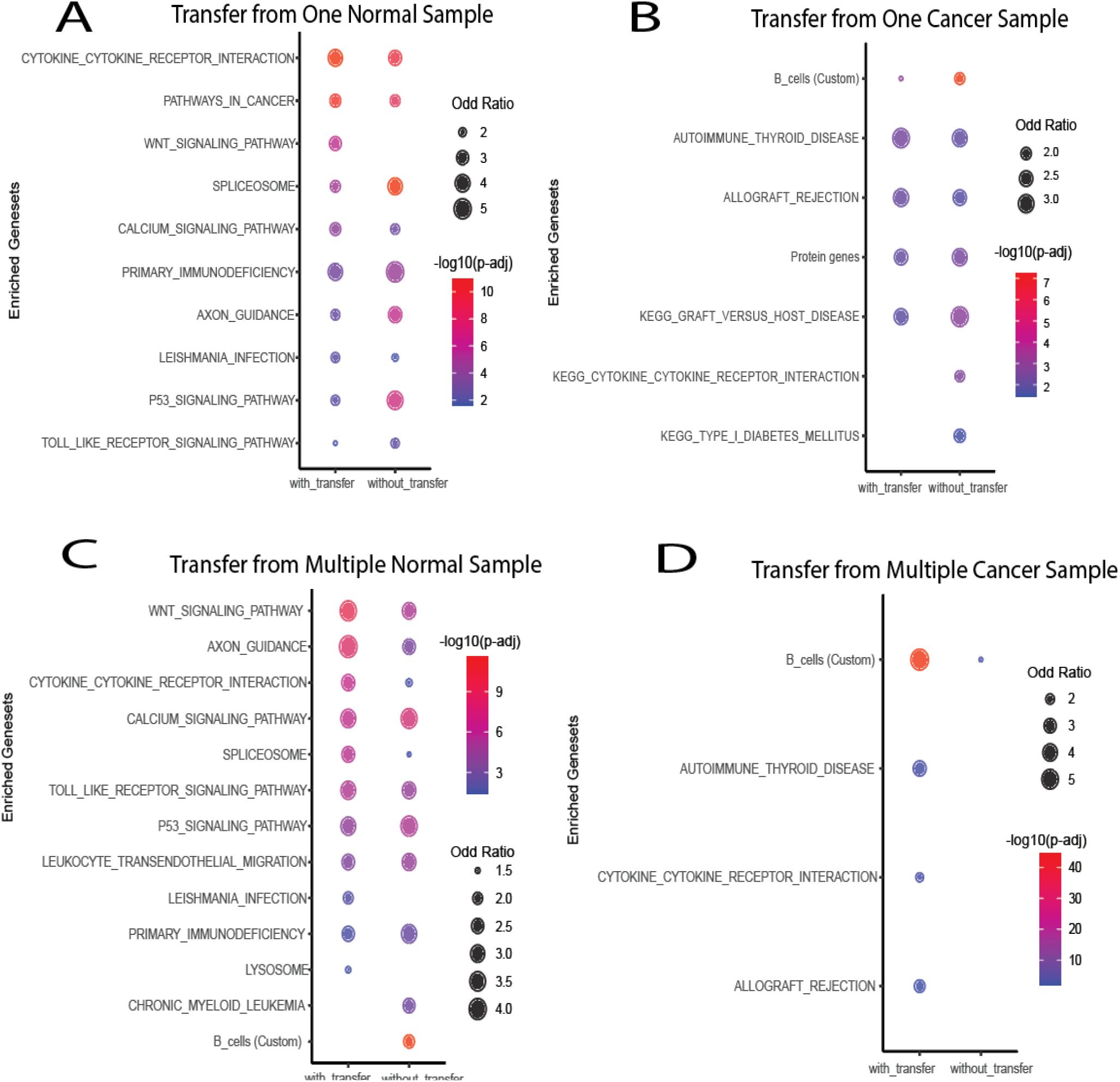
Functional geneset enrichment interpretation of surface protein CD19 using GS-NN and transfer learning for normal and cancer datasets.

For cancer, transfering from C5 GS-NN to C3 actually find less number of enriched genesets comapred to without transfer (Fig. 6B). CD19 prediction PCC perofrmance dropped from 0.56 to 0.43 after transfering. After investigating enriched genesets, we find custom built “B-cells” geneset ranked top with or without trnafering. However, odd ratio of “B-cells” geneset reduced after transfering. This suggests that “B-cells” geneset may have significant contribution to CD19 surface protein prediction in cancer. This should be expected given the performance is worse for cancer.

Next, we invetigate functional differences of with and without tranfer learning using multiple datasets. For normal we are transfering from N1,N2,N4,N5 datasetes to N3. In Section 3.1.3, we report increased average PCC performances after transfer learning with multiple samples for normal. PCC peroformacne of CD19 also imroved from 0.78 to 0.85 after tranfer from four samples. The enrichment results show (Fig. 6C) that ranking of immune related genesets such as ‘WNT_SIGNALING_PATHWAY’,’TOLL_LIKE_RECEPTOR_SIGNALING_PATHWAY’, ‘P53_ SIGNALING_PATHWAY’ increased after tranfer. This may cause GS-NN to learn more robust features for CD19, hence improve PCC performance using present variability from multiple samples.

Lastly, for cancer dataset, tranfering from C1, C2, C4, C5 to C3 improved CD19 predition PCC performance from 0.27 to 0.39. And we find only 4 enriched geneset after tranfering and only 1 without transfering (Fig. 6D). This may be the reason behind low prediction perofemance of CD19. However, comapred to no tranfer we see much improvement after transfering. We also find “B-cells” geneset’s odd ratio incresed significantly with multiple datasets find ‘B-cells (Custom)’ genesets with or without fine tuning with C3. However, odd ratio is increased for ‘B-cells (custom)’ after fine tuning.

## 4. Discussion and Conclusions

We proposed a novel GS-NN and a transfer learning strategy for cross-dataset prediction of cell surface protein levels from scRNA-seq gene expression data. We showed that GS-NN could capture more robust features to enable improved performance of direct cross-dataset prediction especially for cancer. We revealed that improvement is due to the unique geneset-base architecture and geneset level selection process when training a GS-NN model. We further demonstrated the effectiveness of transfer learning and showed that fine-tuning allows further improvements. The unique geneset architecture also facilitates functional interpretation underlying the prediction. The functional analyses of genes with high attribution scores to the prediction revealed that GS-NN and transfer learning make the model focus more on immune-related genesets.

Overall, the importance of this work is three-fold. First, scRNA-seq becomes increasingly accessible, our proposed approach provides a cost-efficient method to augment cell surface protein levels. Secondly, GS-NN is conveniently transferrable across datasets because of the design of the architecture. Lastly, GS-NN facilitates interpretation of biological functions. However, despite the reported improvement by GS-NN over other models, there is ample room for further improvement. Therefore, designing a more robust GS-NN model and transfer learning strategy will be a desirable future research direction.

